# Pupil diameter differentiates expertise in dental radiography visual search

**DOI:** 10.1101/792374

**Authors:** Nora Castner, Tobias Appel, Thérése Eder, Juliane Richter, Katharina Scheiter, Constanze Keutel, Fabian Hüttig, Andrew Duchowski, Enkelejda Kasneci

## Abstract

Expert behavior is characterized by rapid information processing abilities, dependent on more structured schemata in long-term memory designated for their domain-specific tasks. From this understanding, expertise can effectively reduce cognitive load on a domain-specific task. However, certain tasks could still evoke different gradations of load even for an expert, e.g., when having to detect subtle anomalies in dental radiographs. Our aim was to measure pupil diameter response to anomalies of varying levels of difficulty in expert and student dentists’ visual examination of panoramic radiographs. We found that students’ pupil diameter dilated significantly from baseline compared to experts, but anomaly difficulty had no effect on pupillary response. In contrast, experts’ pupil diameter responded to varying levels of anomaly difficulty, where more difficult anomalies evoked greater pupil dilation from baseline. Experts thus showed proportional pupillary response indicative of increasing cognitive load with increasingly difficult anomalies, whereas students showed pupillary response indicative of higher cognitive load for all anomalies when compared to experts.

## Introduction

Mental imagery is a commonly performed task in many contemporary professions, e.g., radiologists and other medical personnel frequently examine medical radiographs to diagnose and treat patients, airport security scan X-rays of luggage for prohibited items, etc. [1]. In such tasks, expertise is derived from domain knowledge and organized schemata that is optimized for a short period of search. Thus understanding the search process, measuring mental workload, and developing computer-based metrics are of fundamental importance.

Of particular interest is estimation of cognitive load during visual search used in demanding real-world tasks. From a human factors perspective, images with complex features can affect performance, especially in visual search, and so selection of measurement techniques to assess human performance is paramount [2]. One especially important factor in performance is attentional cost, where feature complexity has a measurable effect. It is evaluated by one of four general techniques, namely: image processing, objective (performance) evaluation (e.g., task speed and accuracy), subjective evaluation (e.g., self-reported measures, with the oft-used NASA TLX questionnaire being a good example), and eye tracking, which is our focus, and which can be characterized as a physiological measure [3].

Visual search can be split into two stages: search and verification [2], where verification can further be delineated by decision (target/non-target) and confirmation [4]. It is in the verification stage that we primarily expect to find the decision-making aspect of cognition [5–8]. Consequently, we expect that cognitive load measures will manifest significant responses only during the decision-making aspect of the task, when visual search is complete. Just and Carpenter [4] noted that eye fixation data make it possible to distinguish the three stages of visual performance, although their analysis relied on the relation between fixation duration and angular disparity. In our analysis, we focus on fixations during visual search, disregarding saccades, as the moments when we then compute cognitive load from baseline-related pupil diameter difference measures. Other eye movement measures used in visual search and/or cognitive load measurement may also involve fixation/saccade discrimination [2], or metrics based on pupil diameter or microsaccades.

We are particularly interested in examining differences between expert and novice inspectors of dental panoramic radiographs (OPTs), which are information-dense 2D superimpositions of the maxillomandibular region used frequently in all aspects of dental medicine [9]. Due to their heavy reliance on OPTs, dentists undergo professional training and licensing; however, they are still highly susceptible to under-detections and missed information [10–12]. These errors can stem from the technology, e.g. low resolution, high noise, or false positioning, and also interpretation errors [13]. Coupled with concern of patients’ health, accurate interpretation in spite of complex imagery is crucial. Specifically, OPTs have been shown to be less sensitive imagery for certain anomaly types than intraoral(periapical) radiographs, making correct detection more difficult [14, 15]. Therefore, less sensitive imagery of an anomaly can evoke higher gradation of difficulty for accurately interpreting it. Expert dentists are more attuned to the gradation of these anomalies and interpret their image areas accordingly. Therefore, further understanding of both expert and novice OPT examination is necessary for effectively improving the training of medical image interpretation.

### Statement of Contributions

To our knowledge, we are the first to apply differentiable pupillometry to the dental imagery visual search domain. Our contribution is two-fold. First, we show that baseline-related pupil difference, as a measure of cognitive load, is sensitive to experts’ processing of anomalies of varying degree of difficulty. Second, we demonstrate methodological use of a multi-eye tracker classroom for collection of novice eye movement data, which can also serve as a future training classroom, e.g., implementing techniques such as Gaze-Augmented Think Aloud [16].

We start by reviewing domain expertise as a precursor to a review of eye-tracking work in visual search with emphasis on estimation of cognitive processes, then focus on metrics based on pupil diameter to estimate cognitive load.

## Background Characterizing Expertise

Expertise lies in the mind. The theory that expert aptitude develops a more structured long term memory designated for domain-specific tasks [17] offers insight into experts’ faster and more accurate abilities [18]. *Long term working memory*, proposed by Ericsson and Kintsch [17], offers this explanation for how experts seemingly effortlessly handle their domain-specific tasks. Their memory structuring facilitates their ability to maintain working memory at optimal capacity, avoiding overload, which affects productivity and performance.

### Expertise and Memory

Long-term memory (LTM) manages how we automatically engage in familiar activities without much thought (e.g. riding a bicycle, remembering your childhood phone number). Similarly, expertise is dependent on this capacity. However, the mechanism that effectively accesses domain-specific information in LTM is a distinguishing asset to experts. Generally, working memory is understood as temporary storage for processing readily available information [19] and has two prongs: Short-term and long-term working memory. Where the former relates to structuring available for limited capacity, the latter relates to the structuring available to the larger, long-lasting storage and is of more interest in skill learning [17]. This structuring conceptualization explains why experts intuitively handle their domain specific tasks. For instance, chess players employ memory chunking that enables them to quickly recognize favorable positions and movements with less focus on single pieces [20]. Athletes show faster reaction to attentional cues, especially in interceptive sports, (e.g., basketball), indicating more rapid mental processing [21]. Also, medical professionals have been thought to proficiently employ heuristics in their decision making strategies, i.e., visual search of radiographs [22] or diagnostic reasoning [23, 24].

### Skill Acquisition and Cognitive Load

Developing new skills and the related memory structures for a specific discipline rely heavily on the capacity of working memory. According to Just and Carpenter [25], when the working memory capacity is reached, comprehension is inhibited, leading to negative effects on performance. Effective comprehension then relies on *resource allocation* [25]. Optimal resource allocation supports rapid convergence to the most appropriate task-solution. Experts can filter out irrelevant information, which is evident in gaze behavior; they focus more on areas relevant to the task solution and less on areas that are irrelevant to the solution [18]. For instance, expert radiologists have more fixations on anomaly prone areas [26–28] and have shorter time to fixation on an anomaly [22, 29].

Additionally, when the task becomes too difficult, there is more demand on working memory [30]. Sweller points out that the means-to-an-end problem solving strategies that novices employ can overload working memory [31]. For instance, a student using a trial and error approach to an end goal needs to maintain a history of all their wrong answers so far. Each wrong answer then gets added to the stack, taking up working memory capacity. Cognitive load, or more specifically intrinsic cognitive load [32], is the effect of “heavy use of limited cognitive-processing capability” [31]. For more information, see review by Paas and Ayres [33]. High cognitive load has been shown to have negative effects on performance [30] and effective learning in general [34]. Thus, too high cognitive load can hinder the aspect of learning where the memory structures are developed.

Perception of a task as difficult can contribute to higher cognitive load. Although perceived task-difficulty is influenced by acquired knowledge [35], even experts can face challenging problems that could evoke more load on working memory. It has been found that experts employ more efficient reasoning strategies when accurately evaluating clinical case examinations [36]. However, experts that inaccurately evaluated these examinations also employed reasoning strategies similar to novices [36]. Furthermore, inefficient reasoning strategies were also likely to be elicited in experts in more complicated case examinations [37]. Inefficient reasoning strategies brings back the illustration of the stack in working memory being filled with irrelevant information, exhausting capacity and creating cognitive load.

One way to asses levels of cognitive load is the pupillary response [38, 39], where pupil size has been shown to increase as a response to memory resources reaching capacity [40, 41] as well as when the task becomes too difficult [34, 42]. Accordingly, experts have a higher threshold for what is difficult compared to their novice counterparts, which is evident in the pupil response. Therefore, we are interested in expert and novice differences in task difficulty as measured by the pupil diameter. Specifically, expert and novice dentists when interpreting anomalies of varying degree of difficulty in panoramic radiographs. More interesting, our aim is to further understand experts’ perception of difficulty in their domain-specific tasks and whether this affects cognitive load.

## Eye Movement Behavior Reflective of Cognitive Processes

Cognitive processes are evident in the visual search strategy. Generally, visual performance, e.g., during search, has been characterized by metrics derived from the discrimination of fixations and saccades. Fixations are the period when eye movements are relatively still, indicating focus of attention, usually on areas prone to a specific diagnosis [43]. Saccades, the rapid eye movements, are usually made when scanning over irrelevant areas to a specific diagnosis [18]. Kok et al. [44] showed that distinguishable gaze strategies were evident in expert, intermediate, and novice radiologists. Their strategies were affected by *top-down* (context, knowledge-based) or *bottom-up* (salient, noticeable images features) aspects of the task. In other words, top-down based gaze behavior can be representative of the cognitive processing during efficient reasoning. Conversely, less efficient reasoning can be linked to bottom-up based gaze behavior, where attention is spread out over areas deemed salient, regardless of if they are relevant to the diagnosis at hand. Additionally, depending on the anomaly, experts employ a mixture of focal and diffusive, or ambient, gaze strategies [44–46]; however, they are more accurate at determining anomalies than novices and intermediates.

Beyond gaze strategies based on fixations and saccades, other forms of eye movements that have been used to measure aspects of cognition during visual search include their speed and direction [5], microsaccades [47], pupil diameter oscillation [48], and measures related to pupil diameter itself [49]. Generally, most of these measures concentrating on estimation of cognitive load have produced metrics sensitive to the presence or absence of cognitive load. In this paper, we show that the baseline-related pupil dilation, which has been one of the more consistently reliable measures of cognitive load, can also discriminate varying levels of difficulty in experts’ search of radiograph images.

### Pupil Diameter as a Measure of Cognitive Load

Not only does visual search strategy reflect cognitive processes, but pupil diameter has also been shown to be a robust, non-invasive measurement of cognitive load [34, 38–42, 50–55]. Hence, with an increase in task difficulty, the diameter increases, otherwise known as task-evoked pupillary response. Originally, Kahneman and Beatty [50] linked pupil response to attentional differences. Then, the link between attention and capacity was promoted [42]; where higher load on the working memory showed a larger change in pupil dilation. Additionally, pupillary response has been found to be an indicator of learning [34], where pupil diameter decreased with more experience in a task. This understanding of pupil diameter changes has further been employed as a robust cognitive load classifier [39].

Much of the early research in processing capacity and cognitive load have elicited effects from language or number recall tests [40–42]. However, pupil activity correlates to workload during a variety of other tasks (see review by van der Wel et al. [56]). For instance in visual search tasks, more distractors make the paradigm more difficult, affecting the pupil diameter increase [57]. Furthermore, when asked to recall the amount of objects in the stimuli pupil diameter size increased even more [57]. Backs and Walrath [58] found that monochrome displays evoked longer search time and more pupil dilation than colored displays when performing visual search tasks for both object counting and target finding. ^1^ Regarding uncertainty during a search task, an increase in pupil diameter was associated with response time and uncertainty of target selection [59]. Although the effects of learning are still apparent, pupil dilation decreases as an effect of training over time [60].

One of the more important takeaways from the visual search literature is the interplay of selective attention, increasing task demand, and the mental effort evoked. Moreover, this understanding is applicable to medical professionals and the cognitive processes involved during diagnostic interpretation of radiographs, where they are highly exposed to varying difficulties of this task and their student counterparts less so, though they are in the process of learning. Over time, students accumulate more experiences and face more challenging tasks, reducing cognitive load.

### Factors Affecting Pupillary Response

Though it is apparent that pupillary response is a product of cognitive load, other factors have been shown to effect pupil size. For example, changes in luminance in the environment result in the physiological response of constriction or dilation [55]. Age difference has also been shown to affect pupil size differences, where overall pupil size in older adults is smaller than younger adults, though variance between subjects in similar age groups is also quite high [51, 55]. With these factors in mind, studies on pupil diameter and load recommend a task-to-baseline comparison in luminance controlled environments [34, 38–42, 50, 53, 57, 59, 61]. A model was developed that measures pupil dilation during workload that accounts for light changes, where task-related changes are still measurable giving varying lighting conditions [62]. Other factors known to affect pupillary response can be fatigue [63, 64], caffeine or drug consumption [65, 66], and emotion or arousal [67]. Therefore, when measuring pupillary response in relation to cognitive load or mental effort in general, these factors should be controlled in order to avoid such confounds.

## Related Work

When measuring both novice and expert physicians’ performance during clinical multiple choice questions,Szulewski et al. [53] found that novices had a larger pupillary response compared to experts. Also, experts’ pupillary response was not affected by question difficulty or accurate response [53]. Thus, for questions related to field of expertise, trained physicians showed more accurate performance and less cognitive load, whereas novices exhibited greater cognitive load, especially for more difficult questions [53].

Further research in laproscopy found that expert surgeons’ pupil diameter increased as a result of increasing task difficulty during laproscopic procedures [68]. While performing hernia repair surgery, Tien et al. [69] found that junior surgeons had larger pupil sizes than experts and that specific tasks also affected their pupillary response as well. This pupil response due to less experience was corroborated by self-report of task load, where experts experienced less mental demand than juniors [69]. For more references highlighting lower pupillary response as an effect of medical expertise (e.g., surgeons, anesthesiologists, physicians), see Szulewski et al. [70].

Regarding specifically medical image interpretation, Brunyé and colleagues [52] evaluated expert physicians viewing digitized breast biopsies with varying levels of difficulty and their resulting case diagnoses. They found pupil diameter increases as an effect of difficulty in diagnostic decision making, moreso for cases that were accurately diagnosed [52]. They attribute their results to experts’ possible perception of case difficulty during an initial analysis. Therefore, pupil diameter can be indicative of the cognitive processes involved in interpreting medical images and can indicate the level of expertise as well as the degree of difficulty. Brunyé et al. [71] further highlight the prospects that pupillary response in combination with gaze behavior has in understanding uncertainty in medical decision making.

One of the earlier studies that have specifically focused on dental expertise and OPT interpretation found that the degree of image difficulty (obvious, intermediate, and subtle pathologies) had an effect on the gaze behavior for both experts and students [72]. They found that experts were shorter with their total search time as well as time to identify (first fixation) an anomaly compared to novices. However, experts used more fixations and longer fixation durations on difficult images compared to obvious images. Students showed no differences in how often and how long they looked at obvious images or difficult images [72].

Castner and colleagues [73] also found a possible effect of degree of difficulty and how often an expert dentist glances at an anomaly before he or she physically labels it as such. Where certain anomalies were only glanced at once to be accurately labeled, and others needed to be glanced at multiple times to be accurately labeled. Moreover, gaze behavior is indicative of the expertise and the cognitive processes involved in interpreting medical images. Additionally, the degree of difficulty in accurate pathology detection can affect gaze behavior, which can be indicative of the reasoning strategies used. For this reason, we are interested in further understanding the cognitive processes during visual search of dental radiographs. Mainly we wish to know how the degree of pathology difficulty can interrupt the flow of efficient expert reasoning.

With this intention in mind, we looked at expert and novice dentists’ pupillary response while fixating on anomalies of varying difficulty in panoramic radiographs. Not only do these OPTs have multiple anomalies, but also within one OPT, varying difficulties can be present. Therefore, we are not analyzing an overall impression of easy or difficult image. Rather, through the course of the search strategy, we are extracting when they spot an anomaly and extracting the mental processing at that moment. We propose the degree of anomaly interpretation difficulty can be indicated by changes in the pupillary response; where a larger response is more representative of harder to interpret anomalies. We also hypothesize to find a difference in the pupillary response between experts and novices, as established by prior research. However, whether novices are as attuned to anomaly difficulty as their expert counterparts is also of interest to our work.

## Materials and Methods

### Participants

Data collection was performed during summer and winter semesters from 2017 to 2019. Students from semesters six through tenth were recorded during an OPT inspection task. Only the sixth semester students were evaluated three times in each period of data collection due to their curriculum requirement of an OPT interpretation training course. We chose to evaluate the sixth semester students after this course (N_sixthM3_ = 50), since they were more likely to experience cognitive load due to the increase in conceptual knowledge and OPT reading skills from this course. Fig 1 shows the difference in students as well as experts’ pupil diameters over the stimulus duration. The sixth semester students after this training course (“Six M3” in Fig. 4) have higher overall pupil diameter.

**Fig 1.**
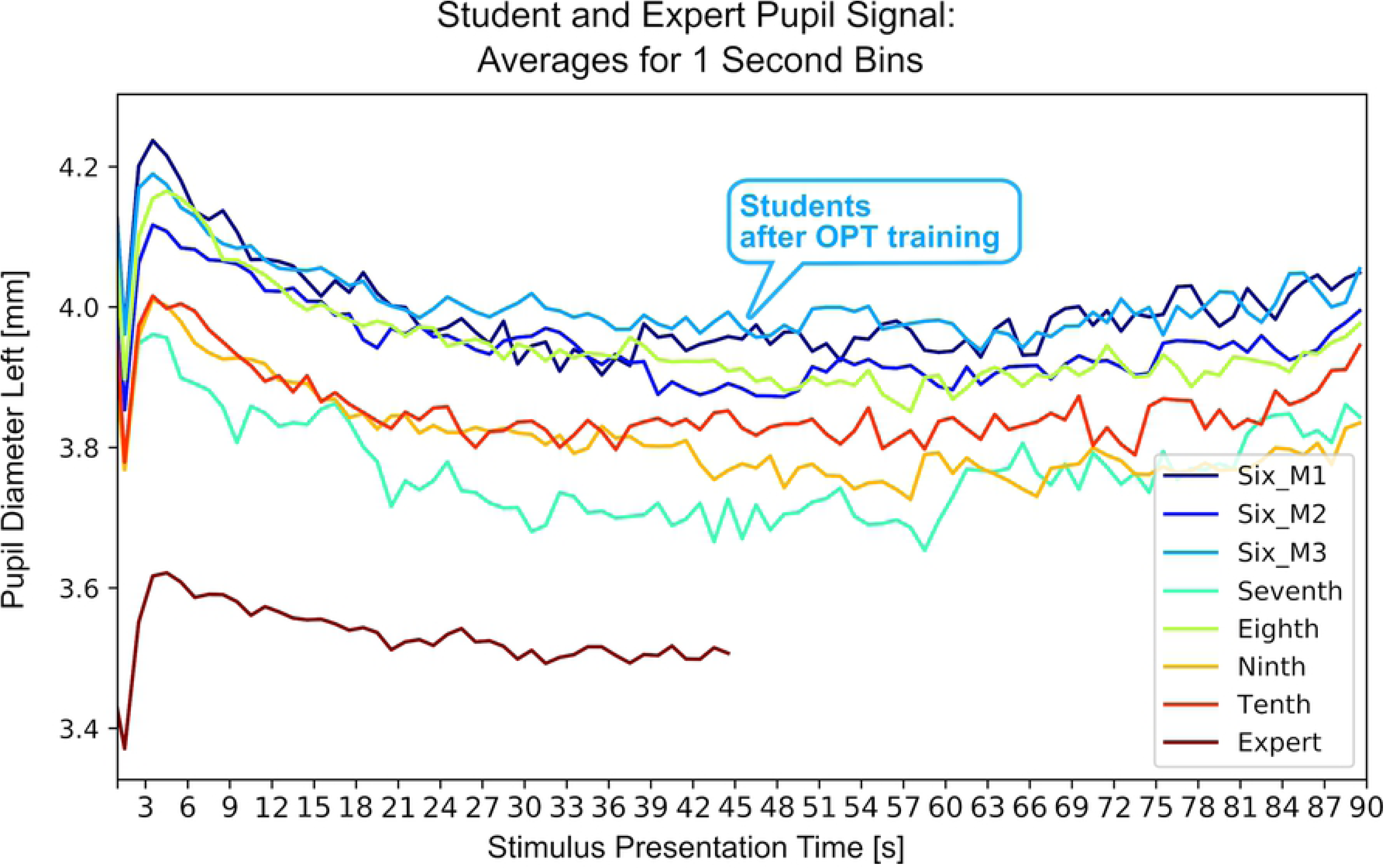
Pupil diameter overtime for students and experts. The raw smoothed pupil diameter overtime of all students collected semesters sixth through tenth and experts. Smoothed raw data is averaged over 1 second bins. The sixth semester students were measured on three separate occasions: Before, during, and at the end of their obligatory training course (as indicated by “M” for measurement). Students had OPT images presented for 90 seconds, whereas experts had the images presented for 45 seconds.

**Fig 2.**
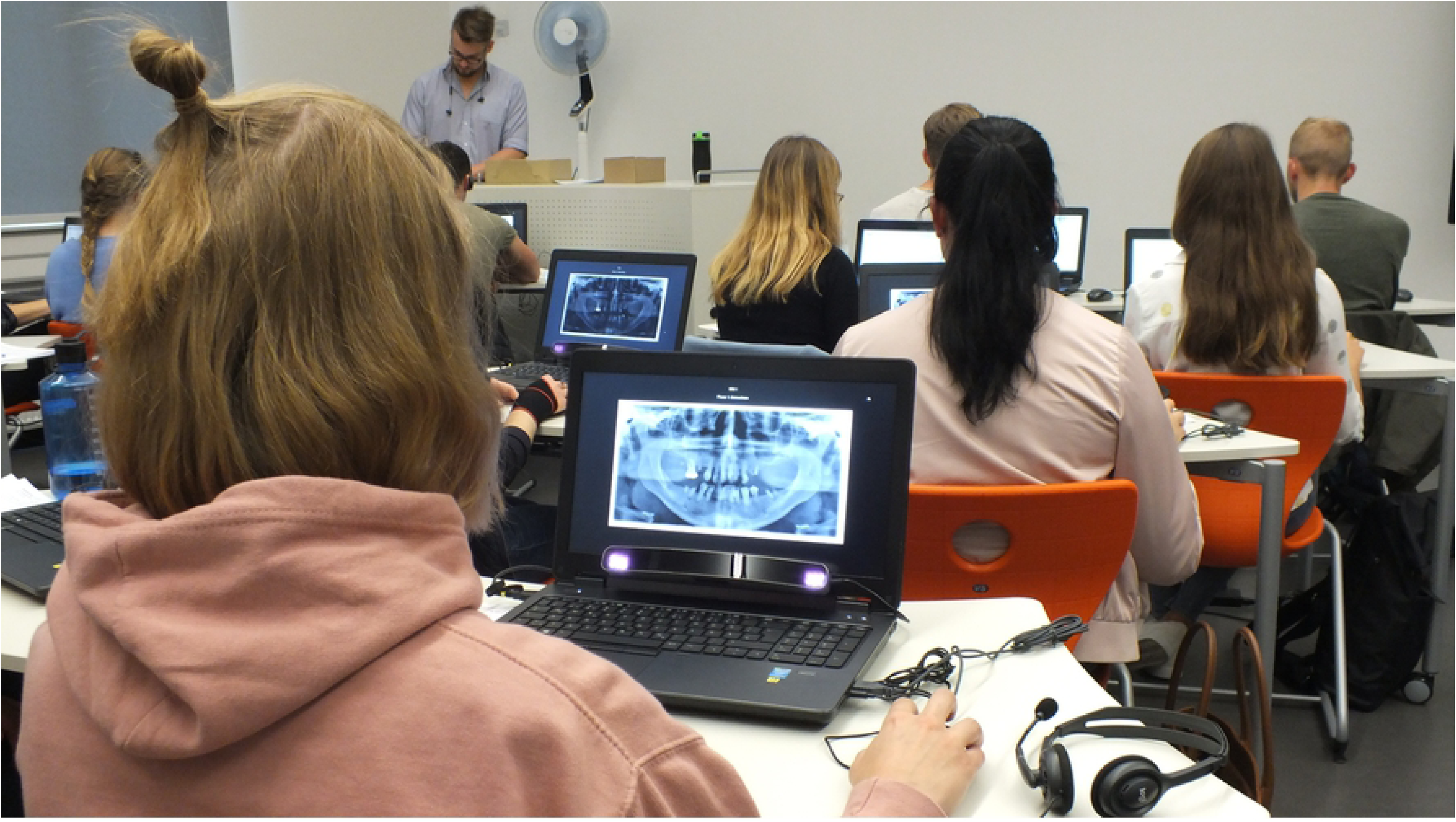
Impressions from the Digital Classroom. This image shows a typical data collection session with students in the digital classroom. It is equipped to handle up to 30 students simultaneously. Each desk offers a laptop with an attached eye tracker, allowing for efficient recording sessions and therefore a large sample size within a short amount of time.

**Fig 3.**
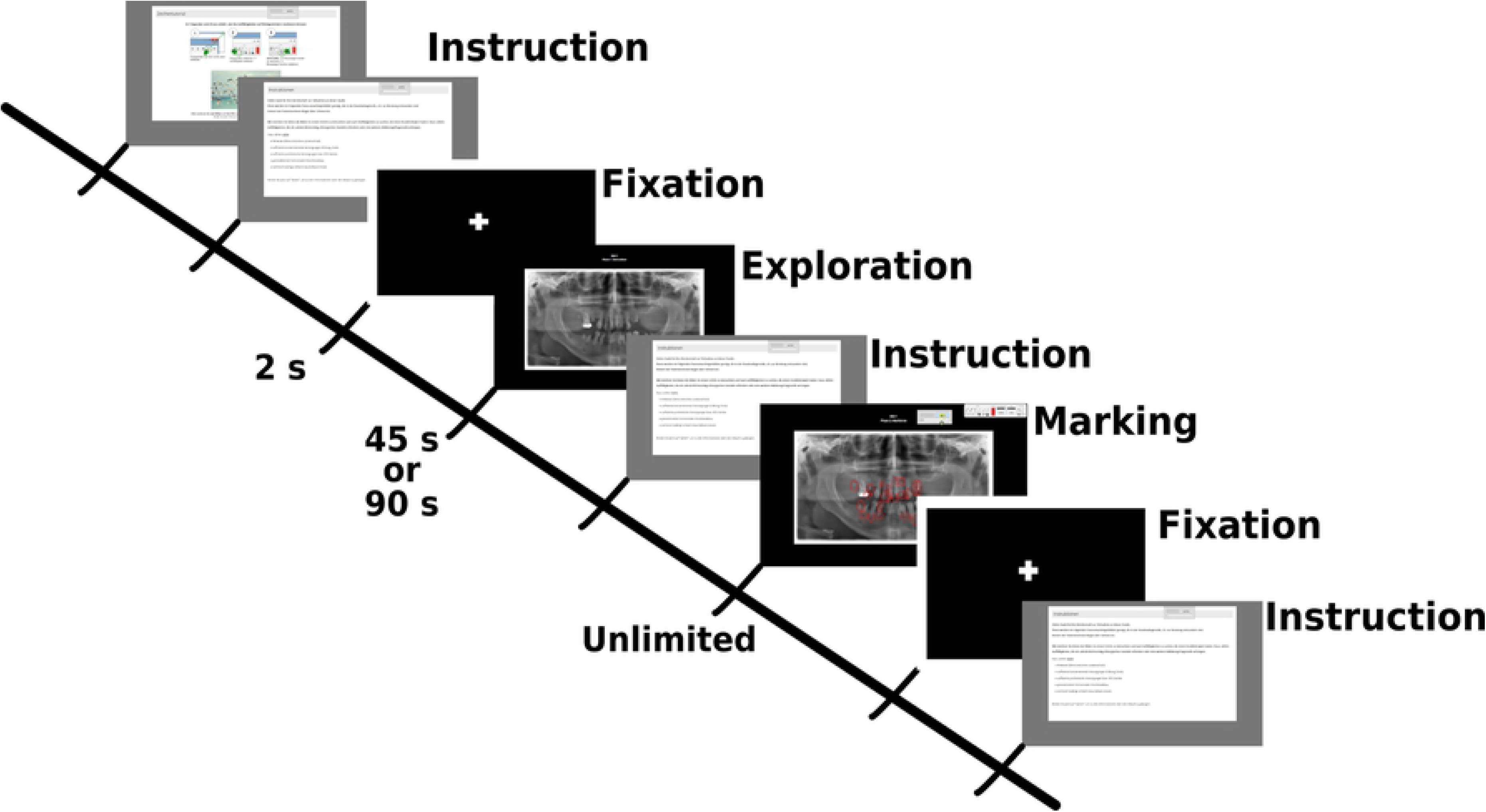
Outline of Experimental Session. Initially, there was a calibration and procedural instructions. Then for each image, there is a fixation cross for baseline data, the exploration phase (45s duration for experts and 90s for students), instructions for the marking phase, and the marking phase (unlimited time). Students received two sets of 10 OPTs with a break in between and experts received one set of 15 OPTs with a break after the first seven.

**Fig 4.**
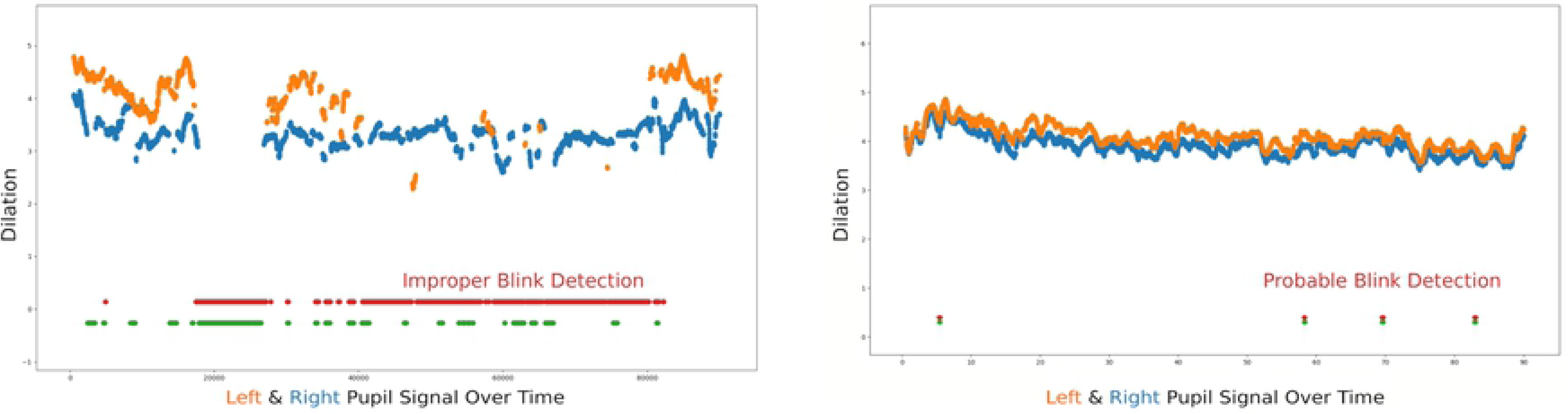
Data Quality Example. (a) Low Data Quality Example (b) High Data Quality Example This graph shows raw pupil signals of the left and right eye over the course of image presentation. Red and green dots in the lower part show when the eye tracker labels the data point as a blink. The particular subject in 4a had a high tracking ratio (98%) for this image, even though there is the possibility that many data samples are missing and incorrectly labeled as blinks. The participant in 4b also has a high tracking ratio, though the data appears to be acceptable with typical blink durations detected and little signal loss.

Table 1 details both the student and expert data. Experts (N_experts_ = 28) from the University clinic volunteered their expertise for the same task that students performed. Experience was defined as professional years working as a dentist and ranged from 1 to 43 years (M_years_ = 9.88). 50% of experts saw between 11 and 30 patients on a given day and the remainder saw less than 10 patients a day. All experts had the necessary qualifications to practice dentistry and or any other dental related specialty: e.g., Prosthodontics, Orthodontics, Endodontics, etc. Due to technical difficulties, eye tracking data was lost for two participants, leaving N_experts_ = 26 participants for the eye tracking analysis.

**Table 1.** Participant Data Overview.

|  | Students 6_M3 | Experts |
| --- | --- | --- |
| $N$ | 50 | 26 |
| $N_{\text{glasses}}$ | 12* | 9 |
| OPTs viewed/person | 20 | 15 |
| Poor Tracking Ratio** | 14.3% | 14.3% |

\*data regarding glasses for one collection is unknown \*\*Percentage of poor data quality. Proportion of valid gaze points less than 80%.

### Environment

Data collection for students took place in a digital classroom (See Fig. 2) equipped with 30 remote eye trackers attached to an HP Z Book Laptop with 17inch fullHD display screen running at full brightness. This special setup allows for data collection of up to 30 participants simultaneously, minimizing the overall time needed for collection. It has the added benefit of synchronized instructions, meaning that all 30 participants present in the classroom receive the same verbal instructions before starting the experiment. Another benefit is that the data of all 30 subjects is collected under the homogenous circumstances: Time of the day, day of the week, specific point in the curriculum, and many other factors that could confound participants’ performance. For this study, verbal instructions were given en masse pertaining to a brief overview of the protocol and an explaination of eye tracking, then individual calibrations were performed with a supervised quality check; students could then run the experiment self-paced.

Data collection for the experts took place in the university hospital so the experts could conveniently participate during work hours. There, the room used for data collection was dedicated for radiograph reading. The same model remote eye tracker was used for expert data collection and was run with the same sampling frequency on a Dell Precision m4800 Laptop with 17inch fullHD display screen running at full brightness.

More important to the current study, both data collection environments had the room illumination levels controlled^2^ with no effects from sunlight or other outdoor light. The standard maintained illuminance for experimental sessions was between 10 to 50 lux: measured with a lux sensor (Gossen Mavo-Max illuminance sensor, MC Technologies, Hannover, Germany). The American Board of Radiology [75] and the Commission of the European Communities advises that environment illumination during radiograph reading should be ambient for the best viewing practices. Ambient light conditions (25–50 lux) optimize contrast perception in radiographs [76–78]. Too bright of an environment can lead to improper luminance transmission [76] from the images, affecting structure discrimination and even effective detection. Therefore, with room illumination controlled, we can evaluate pupillary response independent of environmental illumination changes.

### Laptops

Regarding the screen display and radiograph reading, Goo et al. [77] found effective radiograph reading was not affected by the luminance of the display. Additionally, laptops displays have been found to provide comparable detection performance to other display types [79–81]. However, display standards detailed by multiple medical and radiology commissions are suggested to optimize image quality [75, 82]. For instance, pixel density affects comfortable viewing distances of 30 to 60 cm and a monitor luminance should be at least 200 *cd/m*^2^ to 420 *cd/m*^2^ [75, 83]^3^. Both the laptop models used for the experimental sessions abided by the comission standards. HP Z Book 15 (for students) has screen brightness averages approx. 300*cd/m*^2^ [84]. The Dell Precision m4800 (for experts) averages approx. 380*cd/m*^2^ [85]. While the screen luminance was also controlled and followed the standard protocols for viewing radiographs, the exact effect of the screen brightness on the pupillary response is out of the scope of this work; rather the pupillary response dependent on mental load during these reading task is the focus.

### Eye Tracker

The SMI RED250 remote eye tracker is a commercial eye tracker with 250Hz sampling frequency, and used for gaze data collection. The software included with the eye tracking offers an experiment designer (*Experiment Center*) and event analysis tool (*BeGaze*). Since the eye tracker has a high sampling frequency, both stable (fixations) and rapid (saccadic) eye movements for static stimuli can be measured. Analysis was performed on the raw gaze data output from the eyetracker: *x* and *y* coordinates with timestamps mapped to the screen dimensions. The raw data points also have pupil diameter output in millimeters^4^ Although the data is raw and has not been run through event detection algorithms, raw gaze points are labeled as fixation, saccade, or blink. We evaluated gaze data for the left eye.

Calibration was performed for all participants. Either a 9- or 13-point^5^ calibration was performed in order to accurately map their gaze to the stimuli presentation. A validation also was performed as a quality check to measure the gaze deviation for both eyes from a calibration point. Therefore, if a participant’s validation indicated a high deviation– over one degree– from one or multiple calibration points, the participant performed another calibration. Calibrations were performed prior to the experiments as well as one or two times during the experimental session, depending on how many images were presented.

### Data Collection

The experimental protocol for the students consisted of an initial calibration, task instruction, then two image phases: Interpretation and Marking. The details of the experimental protocol are found in Fig. 3. Prior to the interpretation, a two second fixation cross was presented. Then, an OPT was presented in the interpretation phase for 90 seconds and the participant was instructed to only search for areas indicative of any pathologies in need of further intervention. After the exploration phase, he or she continued with the marking phase. In the marking phase, the same OPT from the exploration phase was shown with the instruction to only mark the anomalies found in the exploration phase with an on screen drawing tool. There was unlimited time for the marking phase, and continued with a button click. This procedure was repeated for all OPTs. In total, the participants view 20 OPTs with a short break after the first ten.

The diagnostic task for the expert group was highly similar to that of the students. However, it was determined that 90 seconds is too long of a duration for the experts, since much of the previous literature has shown experts are faster at scanning radiographs [18, 22, 27, 28, 72, 87–89]. Therefore, the exploration phase was shortened to a duration of 45 seconds. Additionally, due their busy schedules, experts only viewed 15 OPTs, with a short pause after the first seven.

Both students and experts were able to move their head during the experiment, although they were instructed to move their head as little as possible. Further details of one of the student data collections can be found in Castner et al. [90] and expert data collections can be found in Castner et al. [73].

### Pupil Data

#### Gaze Signal

Only gaze data from the interpretation phase was of interest to this work, since gaze data from the marking phase was affected by the use of the screen drawing-tool. Initially, the raw gaze data was examined for signal quality. The eye tracker reports proportion of valid gaze signal to stimulus time as the tracking ratio. Therefore, if a participant’s tracking ratio for an OPT was deemed insufficient–less than 80%–we omit his or her data for this OPT. If overall, a participant has poor tracking ratios for the majority of OPTs he or she viewed (i.e. maximum of three images with acceptable tracking ratios), the total gaze data for that participant was removed. This preprocessing stage can assure that errors (e.g. post-calibration shifts, poor signal due to glasses) in the gaze data are substantially minimized. Table 1 gives the distribution of participants and the percent of datasets excluded due to low tracking ratio (Last row): 199 datasets were initially excluded on the grounds of poor quality data.

#### Blink Information

The tracking ratio does not take into account when the eye tracker detects a blink. Nevertheless, inaccurately detected blinks created an alarming number of cases with acceptable tracking ratios even though there was an inordinate amount of undetected gaze. Fig. 4b shows an example of a participant’s gaze signal (indicated by the pupil diameter value at a given time) for the left and right eye for a 90 second OPT presentation. This participant had a tracking ratio of 98%, but it is apparent that a large portion of the left eye gaze signal– approximately 33.5 seconds– is labeled as a blink by the eye tracker.

Upon investigation, standard and simple implementations for blink detection define a minimum duration threshold that detects a blink if there is no gaze signal for this threshold or longer [43]. The minimum blink duration in the current data set is 70 ms, as corroborated by the SMI manual [86] ^6^. However, it also states that it is not possible for their implementation to distinguish a blink from pupil signal loss [86]. Consequently, the main issue stems from the apparent lack of a maximum blink duration threshold. The question becomes, did the student in Fig. 4a close his or her eye –Pirate style– for almost half of the stimulus, or is this simply a situation where the left eye was not detected for more than half the time?

Extra criteria was necessary to further detect and exclude datasets with pupil signal loss mislabeled as a blink. We overestimated the threshold for atypical blink durations, setting this value to 5000 ms, to account for situations where a participant could possibly be rubbing his or her eye/s or possibly even closing the eye shortly. This threshold could then optimally leave an acceptable amount of pupil data for the entire stimulus presentation (90 or 45 seconds). Since baseline data was the two second fixation cross presented directly before each stimulus, we set the threshold blink duration to 500 ms and added an extra criteria of a minimum 200 pupil samples to effectively extract enough samples for an acceptable pupil diameter baseline. Therefore, an initial quality check was low tracking ratio exclusion. Then, the second data quality check removed data sets if blink durations were atypical. These datasets were excluded from the final analysis, leaving 570 datasets from 72 participants (48 students, 24 experts).

#### Pupil Diameter

Data analysis was done for the left eye. As previously mentioned, the raw gaze signal is divided into the labels blink, fixation, or saccade. Thus, we can determine when the gaze signal is indicative of fixation-like behavior and saccade-like behavior. For further signal processing, we removed gaze coordinates and pupil data for the raw data points labeled as saccades (since visual input is not perceived during rapid eye movements [43]). Data points with a pupil diameter of zero or labeled as a blink were also removed. Additionally, data points 100 ms before and after blinks were removed, due to pupil size distortions from partial eye-lid occlusion. Lastly, the first and last 125 data points in the stimulus presentation were removed due to stimulus flickering. [91–93] The remaining data was smoothed with a third order low-pass Butterworth filter with a 2Hz cutoff as illustrated by the purple data points in Fig. 5.

**Fig 5.**
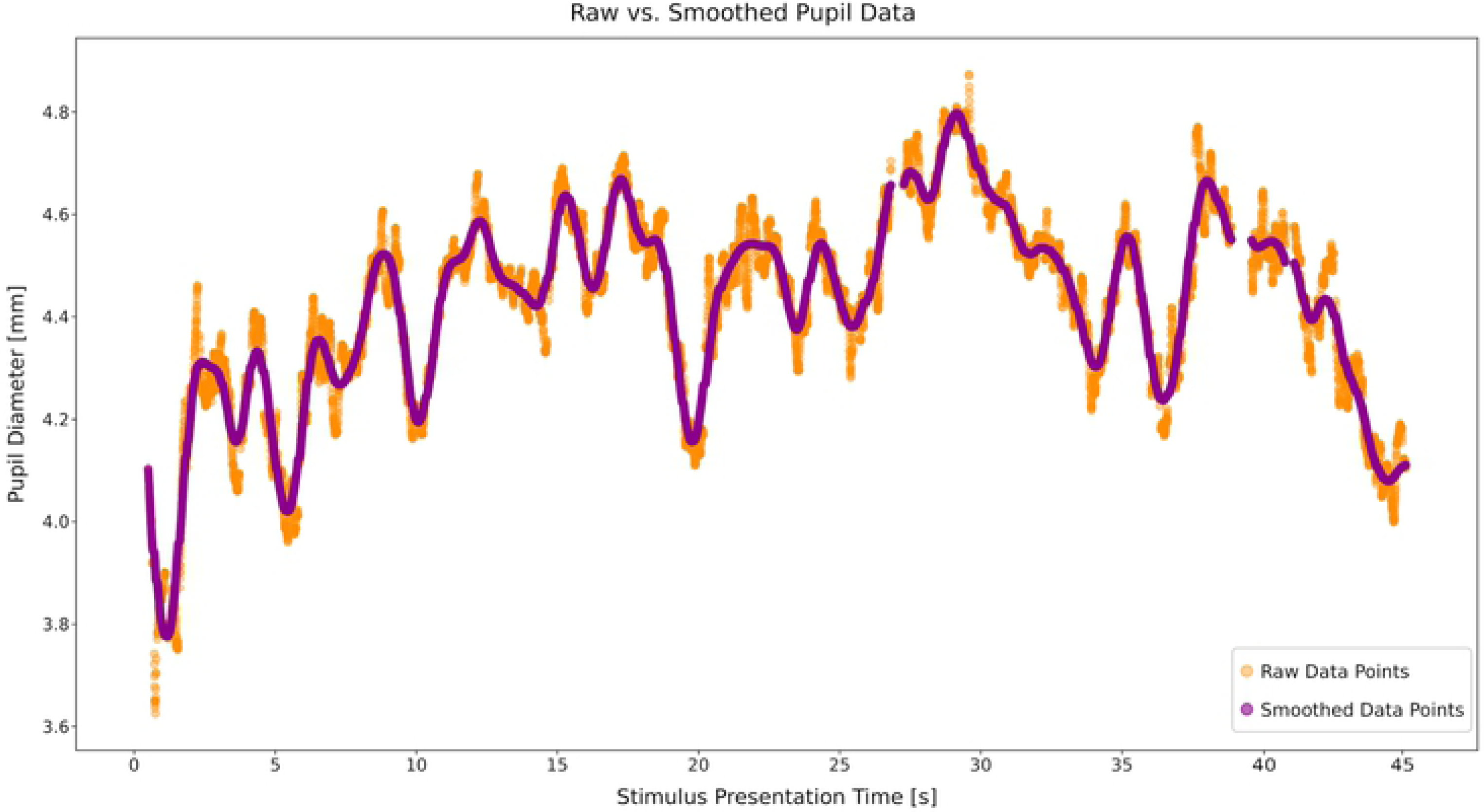
Smoothed Pupil Signal. Raw signal from the left eye (orange) and the smoothed signal (purple) with a butterworth filter with 2Hz cuttoff.

### Pre-Determined Ground Truths

The sixth semester students evaluated in this study (M3: post training course) viewed 20 OPTs and the experts evaluated viewed 15 of the same OPTs. The OPTs were chosen from the university clinic database by the two expert dentists involved in this research project, and were determined to have no artifacts and technological errors. Both dentists independently examined the OPTs and the patient work-ups and further consolidated together to determine groundtruths for each image. Two OPTS were negative (no anomalies) controls.

#### Anomaly Ground Truths

Additionally, the level of difficulty for each anomaly was pre-determined. Fig. 6 shows four OPT images viewed in the experiment. Anomalies are illustrated in green, yellow, and red, and represent easy, medium, and difficult respectively^7^. For example, the green anomalies in Fig. 6 (A) are dental cyst (1) and insufficient root canal fillings. (2a,b) in Fig. 6 (C) are an example of elongated lower molars due to missing antagonists. The yellow anomalies in Fig. 6 (B) are irregular forms of the madibular condyle (1,3) and (2) is an apical translucency indicative of inflammation due to a contagious (bacterially colonized) root canal filling. The red anomalies in this image are approximal caries (4) and a maxillary sinus mass. Anomalies indicated by the white dashed circles were determined as ambiguous, e.g. the nature of their difficulty and or pathology is unclear. For example, in Fig. 6(B)(7,8) are impacted wisdom teeth, though it is uncertain whether this will become a problem for the patient and therefore is regarded as potentially pathologic. (6) is an apical translucency at the mesial root apex and it is unclear whether it is indicative of an inflammation. Therefore, they were kept in this analysis even though the nature of their difficulty is unclear.

**Fig 6.**
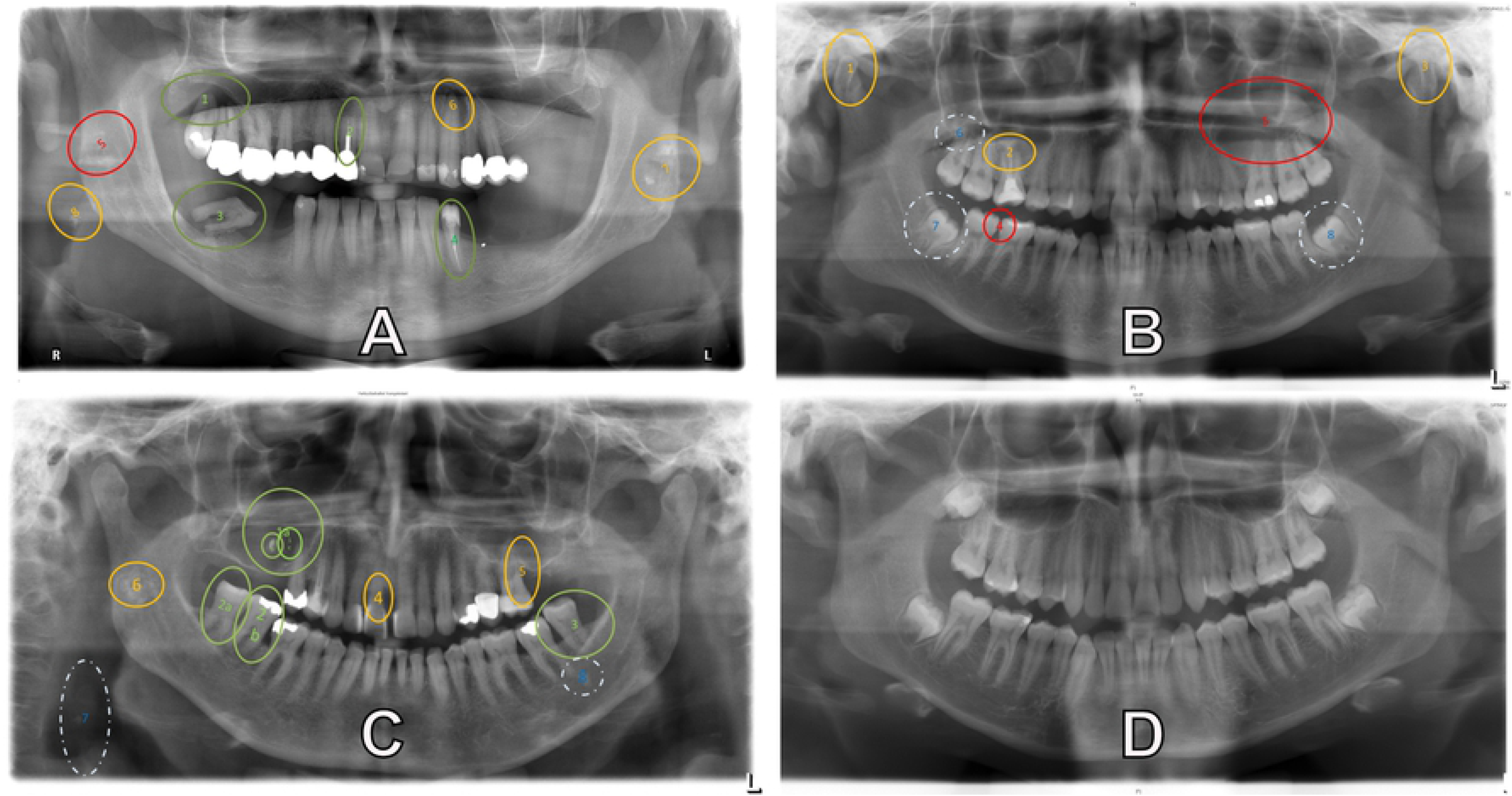
OPTs with Pre-determined Ground Truth. Example of the OPTs used in the experiment. Pre-determined ground truths are indicated by the ellipses and their colors indicate the level of difficulty each anomaly is: Green(least difficult), yellow (intermediary), red (most difficult) and white (nature of difficulty unclear). Image (D) is a negative control image with no anomalies.

#### Anomaly Maps

We created maps for the 15 OPTs evaluated (See Fig. 7) using Matlab 2018. As input, all OPTs were loaded as .png files with their respective anomalies– all colored red. Thresholding for red values was performed to automatically get the pixel coordinates of the ellipse edges. Then, the ellipses were filled with the `poly2mask()` function. Anomalies automatically extracted from this process were double checked for overlapping and had their boundaries corrected. Certain anomalies inside another and that were highly similar in nature, such as (2a,b) in Fig. 6(C), were grouped together as one anomaly. Other anomalies too close together and too different in pathology, such as (3,8) in Fig. 6(C), were excluded from the analysis, due to possible spatial accuracy errors in the gaze. Similarly, anomalies that were denoted by too small of an ellipse were padded to have a larger pixel area,e.g. (4) in Fig. 6(B), to account for an spatial accuracy errors in the gaze. Each segmented anomaly is given a distinguishing integer for its respective pixels. Raw gaze points from the left eye are then mapped to the map and gaze coordinates receive the corresponding integer value.

**Fig 7.**
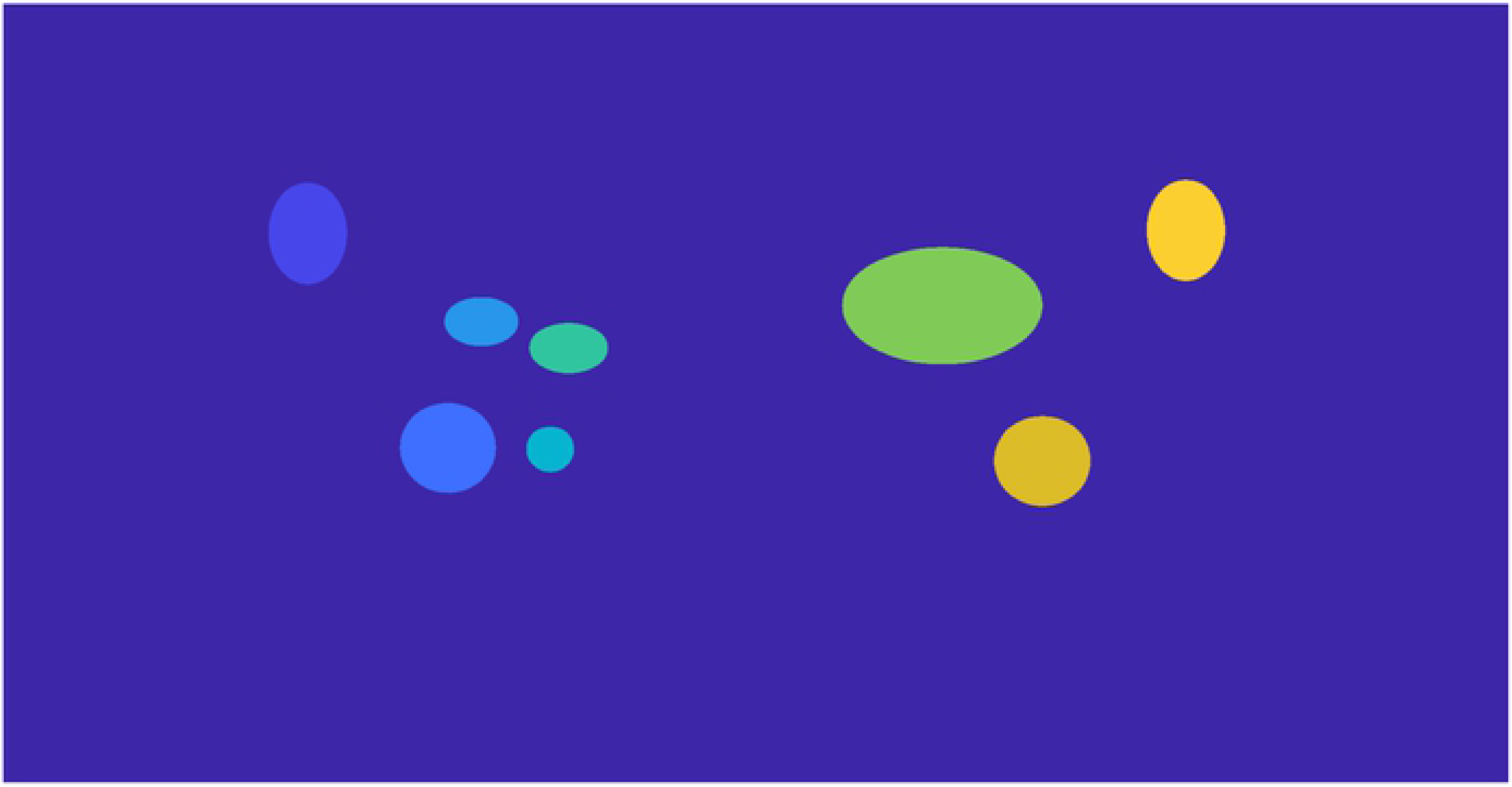
Map of Ground Truths. For image (B) in Fig. 6. Each anomaly is segmented and given a distinguishing interger. Raw gaze points from the left eye are then mapped to the map and gaze coordinates receive the corresponding value. These distinguishing values are further linked to the pre-determined anomaly difficulty in order to get a count of how many raw gaze hits landed on each anomaly type.

### Gaze Mapping to Anomaly

For both students and experts, we plotted the raw gaze points that landed in each anomaly and extracted its level of difficulty. For simplicity, we will refer to them as gaze hits. For all hits on an anomaly for a participant, we calculated the median pupil diameter. The median pupil diameter for each anomaly was then subtracted from the respective baseline data for that image. Therefore, the difference from baseline could be indicative of diameter increase (positive value) or diameter decrease (negative value) compared to baseline.

With the gaze hits on anomalies of varying difficulties, we can evaluate the pupillary response of both experts and students during anomaly fixations. The pupillary response, as measured by change from baseline, can then provide insight into the mental/cognitive load both groups are undergoing while interpreting the anomalies.

## Results

### Overall Change from Baseline

Independent of gaze on anomaly behavior, we looked at participants’ median pupil diameter for each image compared to baseline median pupil diameters. We favored the median over the mean, because it has greater robustness towards noise and outliers. Fig. 8 shows the average of the median pupillary response from baseline for both students and experts. Overall, students (M = 0.314, SD = 0.315) had a larger change from baseline than experts (M = 0.057, SD = 0.353 : *t*(568) = −8.824, *p* < 0.001).

**Fig 8.**
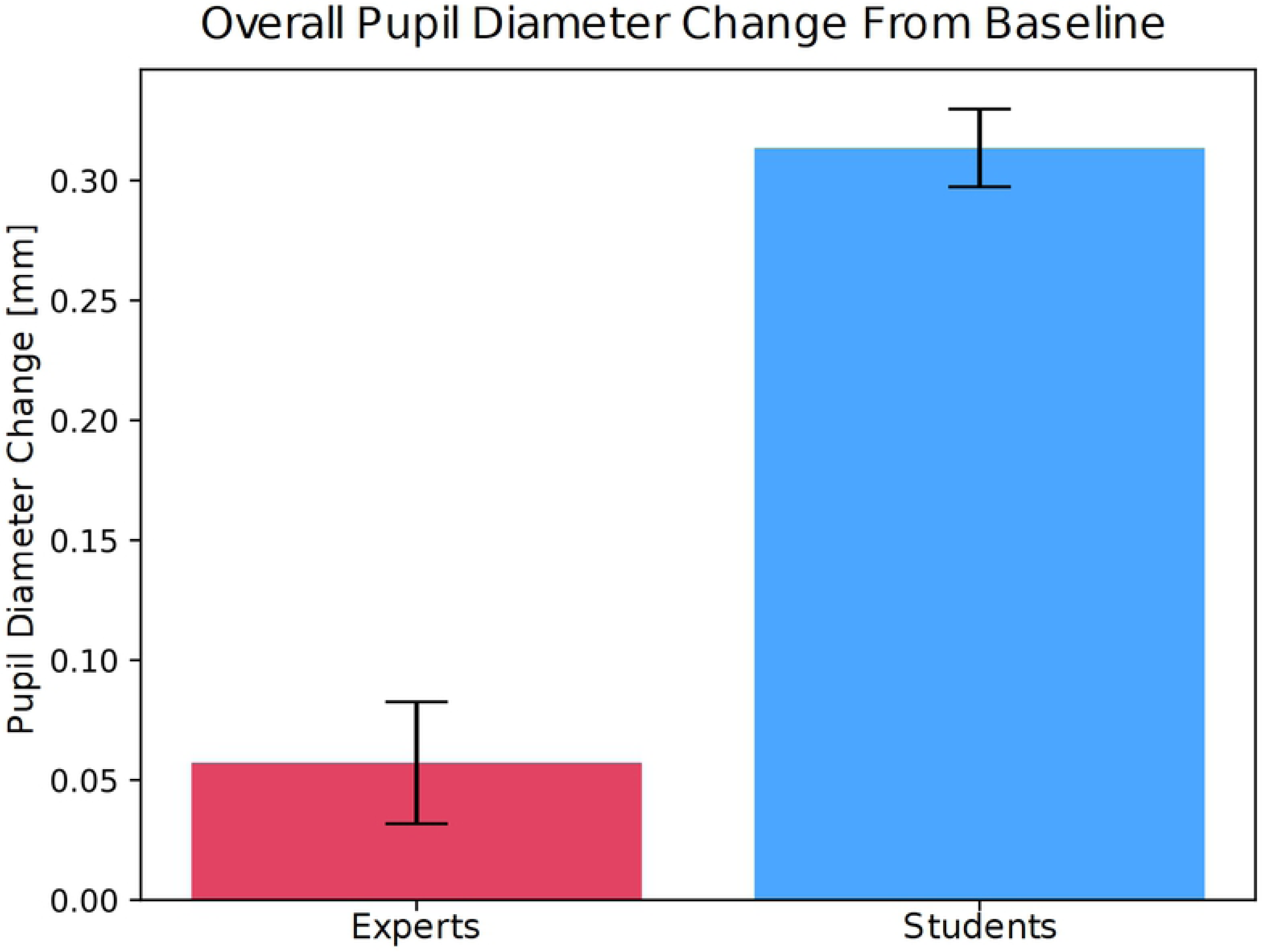
Median Pupil Change From Baseline. The average pupillary response from baseline for students (blue bar) and experts (red bar) during visual search of the whole OPT, regardless of gaze on anomalies. Standard errors for both groups are indicated by the black lines. Students had significantly larger pupillary response from baseline while visually inspecting the OPTs.

### Gaze on Anomalies

To evaluate whether anomaly difficulty had an effect on student and expert pupillary response, we ran a one-way AVOVA for both experts and students. There were no significant effects of anomaly difficulty on student pupillary response (*F* (3, 930) = 1.33, *p* = 0.26). However, there were significant effects of anomaly difficulty on expert pupillary response (*F* (3, 458) = 4.39, *p* = 0.0046). Experts had small pupil diameter change from baseline (M_Expert_ = 0.246, SD_Expert_ = 0.370) except when gazing at more difficult anomalies. Students had large changes from baseline (M_Student_ = 0.367, SD_Student_ = 0.306) for all anomaly difficulties; Fig 9 details this behavior.

**Fig 9.**
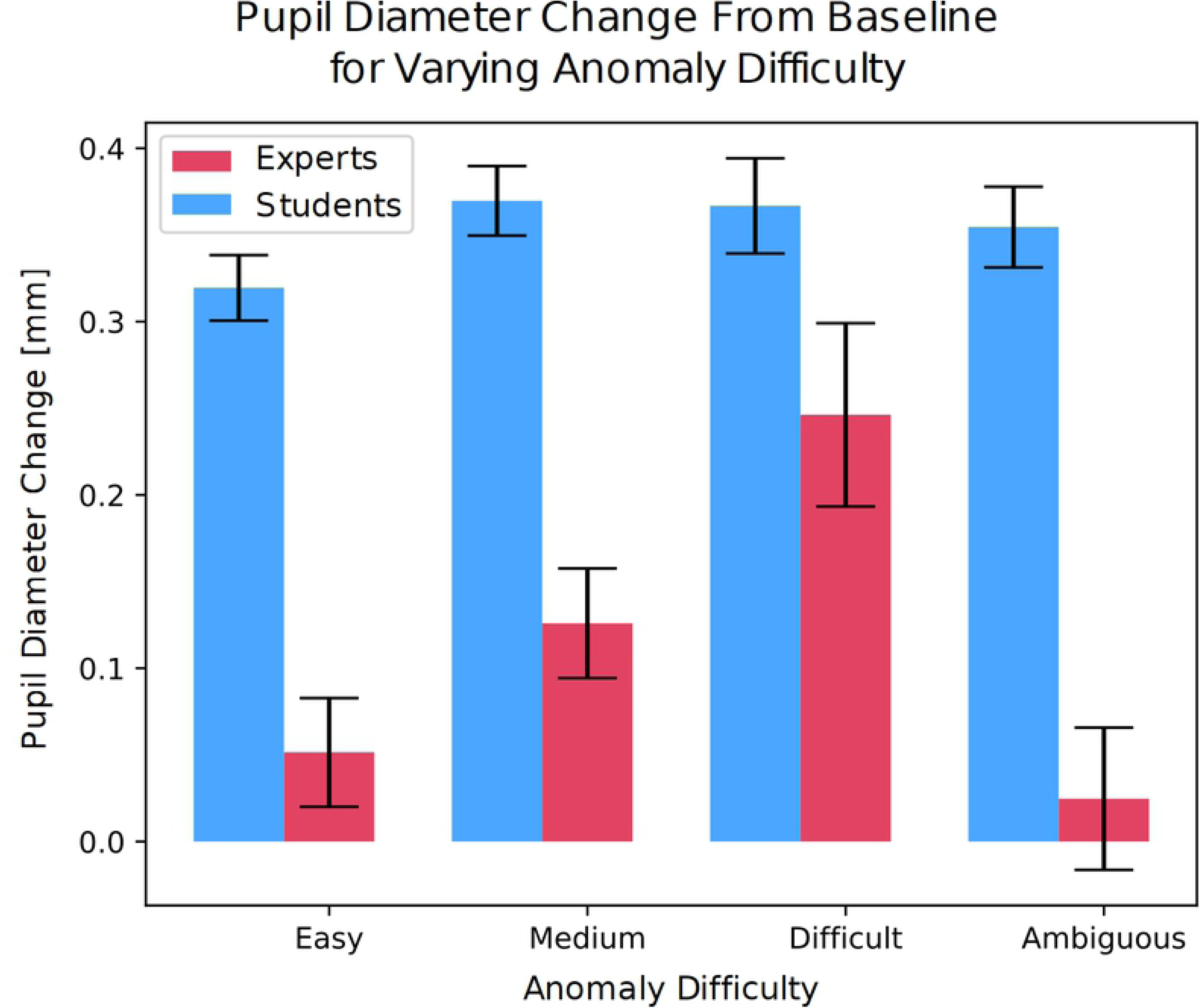
Median Pupil Change From Baseline for Gaze on Anomalies. The median pupil diameter change from baseline for students (blue bars) and experts (red bars) when gazing on anomalies of varying difficulty. Standard errors are indicated in black. Students had larger pupillary response from baseline compared to experts but this effect was homogeneous for the differing anomalies. Whereas experts showed a pupillary response behavior as an effect of increasing difficulty. Though it is unclear as to the nature of the anomalies pre-defined as ambiguous.

A 2 × 4 factor ANOVA to test for interactions found significant effects for expertise (*F* (1, 1388) = 161.68, *p* < 0.001), anomaly (*F* (3, 1388) = 3.87, *p* = 0.009), and the interaction between expertise and anomaly difficulty (*F* (3, 1388) = 2.76, *p* = 0.041). Post hoc analysis with Bonferroni correction for anomaly difficulty on the expert data revealed significant differences for the more difficult anomalies (M = 0.246, SD = 0.370) compared to least difficult (M = 0.0514, SD = 0.396, *t*(207) = −3.0582, *p* = 0.002) and ambiguous (*t*(150) = −0.3988, *p* = −0.044) Meaning, experts had the highest pupil size change from baseline for more difficult anomalies, especially compared to least difficult and ambiguous anomalies, which both had less pupil diameter change from baseline.

## Discussion

We measured pupil diameter change from baseline when gazing on anomalies of varying difficulty during visual search of dental panoramic radiographs. We found that the gradation of anomalies in these images had an effect on expert pupillary response. Anomaly gradation did not have an effect on student pupillary response.

Students showed larger and more homogenous pupil size change from baseline for all anomaly gradations compared to experts. Thus for students, pupillary response was independent of whether an anomaly was easy or difficult to interpret. This effect was also found during visual inspection of the whole image (Fig. 8). Students showed significantly larger pupillary response than experts, which has been supported by the previous literature [52, 53, 69–71, 94]. This response has also been indicative of higher cognitive load [25, 40–42, 50, 53, 92]. For instance, Tien et al. [69] found that novices have more higher memory load compared to an expert performing the same task. This behavior can be likened to students’ lack of conceptual knowledge and experience producing them to “think harder” [95, 96] to interpret these images.

The more interesting takeaway from this work is the lack of influence of anomaly gradation on student cognitive processing. One would imagine that even the most pronounced of anomalies would make the recognition process easier. However, the pupillary response indicates that, regardless of how conspicuous, the level of mental workload remains constant. The large pupil size could be reflective of learning during the task, where students are developing the proper memory structures as theorized by Ericsson and Kintsch [17]and Sweller [31]. Additionally, it could reflect that they have not yet developed the conceptual knowledge to quickly recognize the image features indicative the specific anomalies or how to interpret their underlying patholgies. Even for easy anomalies, they may be unsure of whether they accurately interpreted or not.

Therefore, pupillary response while focusing on anomalies during visual search of OPTs suggests students employ similary cognitive strategies for differing anomaly gradations. Patel et al. [36] found this similar behavior when novices interpreted clinical case examinations. Furthermore, previous reseearch has found systematic gaze strategies were similary present in students searching medical images [22, 28, 44]. Systematic search has also been shown to affect larger pupil dilation [97]. Systematic search evokes more load on the working memory, however, this is what the students are generally being trainied to perform, when they first get exposed to these images [89, 98].

Conversely, experts showed a strong pupillary response to anomaly gradation. Where the least difficult to interpret anomalies showed less change from baseline, then the intermediary anomalies, and finally the largest response was for the most difficult anomalies (Fig. 9). Meaning, as the gradation of difficulty increases so does the pupillary response. This behavior, however, was not evident for the ambiguous anomalies, which showed the smallest response change from baseline. This behavior effect may lie in the nature of the uncertainty of these anomalies. As determined by the two experts involved in the project, this category was a mixture of potential areas that may or may not have included an anomaly: Or even an anomaly, but with no cause for alarm. Therfore, it is uncertain how difficult, easy, or even existing these anomalies were.

Nevertheless, when expert dentists perform a visual inspection of an OPT, they gaze in many areas that potentially have a multitude of differing pathologies or even positional and summation errors. Depending on the gradation of the area they are focusing on, proper interpretation may need to evoke differing processing strategies. In general, as task difficulty increases, so does the workload [68] and correspondingly, the pupil dilation [30, 42, 48, 99]. Patel et al. [37] found more cognitive load in physicians examining more complicated case examinations. Duchowski et al. [47] also showed increased cognitive load during decision-making of increasingly difficult abstract stimuli, but did so using microsaccade rate. Chi et al. also found that experts can more accurately determine how difficult a problem is [100].

Gaze behavior in expert dentists was also shown to change with difficult images [72]. Castner and colleagues [73] also found that different image types evoked either more or less fixations in order to accurately detect anomalies. The current work went one step further and found changes within the visual search of an OPT in contrast to the overall response to interpretation of such an image. In visual search, employing a *top-down* strategy means that someone uses his or her acquired knowledge and understanding of the current problem to focus on the relevant aspects of an image to effectively process it [26, 98, 98]. Moreover, prior knowledge to a problem has been shown to reduce cognitive load [31, 33, 40, 53]. An expert generally knows in what areas of the OPT they are prevalent and how they are illustrated in the image features. Therefore, from these top-down effects, an expert can quickly recognize an image feature as a specific anomaly. In contrast to overall visual inspection, were we found that experts showed low average change from baseline. When inspecting specific areas, pupil dilation fluctuation can be indicative to changes in workload even for experts. Although, experts have a higher threshold for perceived difficulty than students, it is assumed that they still experience tasks or subtasks they perceive as difficult or can be uncertain about.

However, if all anomalies and their pathologies were equally prevalent and salient in OPTs or any other medical image types, experts could effortlessly detect the vast array of issues with 100 percent accuracy. Also in this case, training of accurate detection would increase solely from more exposure. Naturally, interpretation of medical images is not this simple and certain image or pathology features can avert the true diagnosis. Experts are more robust at determining more difficult or subtle anomalies [12, 28, 72, 89, 101]. Although harder to detect anomalies evoke behavior indicative of task-difficulty [34, 42]. More subtle anomalies evoked behavior that is likely of more thorough inspection.

Experts, though reknowned for their streamlined processing abilities, are able to selectively allocate their attention to relevant information and is evident in the pupillary response. However, selective attention coupled with focus on an area perceived as challenging can increase the pupil dilation even further as we found in our investigation. Similar to students, albeit perceived to a lesser extent and only for difficult anomalies, is the effect of uncertainty on the pupil size when looking at these specific image features.

## Conclusion

In short, we found evidence of workload in experts as well as differences between expert and novice workload during visual inspection of dental OPTs. However, it should be noted that there were age differences between the two groups. Due to the sensitivity of the expert demographic data, we did not record their ages; but we can expect them to be older than their student counterparts. Age has been found to have an effect on the average pupil size [51, 55]. For this reason, we measured a change from baseline. Additionally, Van Gerven et al. [54] found that pupillary response to workload in older adults is not as pronounced as in younger adults. However, their population was adults in their late sixties and early seventies compared to adults in their early twenties [54]. Though we cannot say exactly how old our expert population was, they were all still working in the clinic and therefore more than likely to be younger than early seventies. Also, their years of experience in the clinic (average of 10 years) suggests they were more middle aged (30 to 45 years old). Further research is needed to better address this limitation control for possible age difference effects on pupillary response.

Another limitation to this work could be the technical problem associated with the eye tracker data collection. We removed data sets determined as poor quality; however, spatial resolution errors can accumulate within an experimental session if a participant moves too much. Then, the gaze appears to have a shifted offset, which would affect precision in determining if a participant looked at an anomaly. To control for this error, we increased the areas of smaller ground-truth anomalies and excluded anomalies that were too close and too different in nature. The total gaze hits on each type of anomaly were not evenly distributed, with more gaze hits on easier and intermediary anomalies. Students used more total gaze hits due to longer OPTs persentation time, but the distributions were highly similar to experts. Future research can further untangle the differences in gaze hits on easier and difficult anomalies, while controlling for presentation time differences.

Although a majority of expert studies have established that experts are more robust at accurately solving their domain-specific tasks than their student counterparts [17, 18, 26, 102], pupillary response during anomaly inspection in connection to detection performance is also of interest for furture work. It would be interesting to see whether pupil diameter may be indicative of not only anomaly difficulty but also accurate detection of difficult anomalies.

The temporal scanpath information is also an interesting direction for future research, where systematic search in students and its effect on workload and pupillary response. For example, how often do “look backs” on anomaly areas occur and does the pupil dilation increase with each look back. Also, whether easy or more conspicuous anomalies are viewed at first and how the pupillary response in students incorporates this initial information. Following up on the understanding that systematic search produces more memory load as measured by pupil dilation [97], would also be interesting to replicate with temporal information from our findings.

## Supporting information

**Table 2.** Supplementary Table. Raw Gaze Count on Anomaly. **Supplementary Table. Table of Expert and Student Gaze Counts**. shows the gaze hits on each anomaly type for both students and experts. For both levels of expertise, the least difficult and intermediate have the most gaze hits. The following are the ambiguous and the most difficult anomalies. Students had overall more gaze hits than experts; however, this may be attributed to the 90 second viewing time they had in comparison to the 45 second viewing time that the experts had.

| Anomaly Type | Less Difficult | Intermediate | More Difficult | Ambiguous |
| --- | --- | --- | --- | --- |
| Total | 471 | 448 | 173 | 304 |
| Student | 312 | 296 | 124 | 202 |
| Expert | 159 | 152 | 49 | 102 |

## Footnotes

1 They could not rule out luminance differences as a possible confound.

2 Illuminance is the amount of light on a given space. Luminance is light reflected off a surface [74].

3 Depending on nature of radiograph, e.g. Mammography, CT.

4 Millimeters extrapolated from pupil height and width dimensions in pixels [86].

5 The first data collection of the sixth semester students was done with 13 points. However, the other data collections were done with 9 points. The sixth semester students and experts analyzed for this work both performed 9-point calibrations.

6 In general, other parameters that can further customize blink detections can be pupil diameter change and velocity [43]

7 This classification was set up in a blinded review and the consent process of two senior dentists (6th and 7th authors).

